# Glycosyltransferases regulate the expression of Golgi phosphoprotein 3 (GOLPH3)

**DOI:** 10.64898/2025.12.31.695843

**Authors:** Natalia Martínez-Koteski, Fernando M. Ruggiero, Sofia Rasino, Pablo H. H. Lopez, Gerardo D. Fidelio, A. Alejandro Vilcaes, Natali L. Chanaday

**Author notes:** Correspondence (A.A.V.), (N.L.C.).

## Abstract

Glycosphingolipid glycosyltransferases (GGTs) can organize as multienzyme complexes localized along the Golgi complex. However, the influence of the relative presence of GGTs on the localization of their clients is unclear. Here, we determine that expression of certain full-length GGTs increases the levels of Golgi phosphoprotein 3 (GOLPH3), an adaptor oncoprotein involved in Golgi trafficking and organization. Furthermore, we demonstrate that expression of the N-terminal domain of GGTs, which lacks the catalytic domain, is sufficient to achieve this regulation on GOLPH3 in a cell type-dependent manner. We also identify the N-terminal domain of β4GalT-VI GGT as an inhibitor of GOLPH3 expression and thus a potential therapeutic application, since GOLPH3 overexpression is associated with progression and poor prognosis of multiple tumor types. Our data further suggest that the cytoplasmic tail of β4GalT-VI N-terminal domain interferes with the ability of GOLPH3 to interact with phosphatidylinositol 4-phosphate, which consequently reduces the levels of GOLPH3, thereby impairing its function in the acquisition of mesenchymal features.

## Introduction

Glycosyltransferases play a crucial role in cell physiology, contributing to glycosylation of proteins and lipids. Their abnormal expression and/or activity are associated with several human diseases and disorders (Cumin et al., 2022; Lopez & Schnaar, 2009; Varki et al., 2022). The canonical function of glycosyltransferases is to catalyze the transfer of a monosaccharide from an activated donor molecule to a specific acceptor molecule, forming a glycosidic bond. With few exceptions, the enzymatic activity of glycosphingolipid glycosyltransferases (referred as GGTs) is mainly carried out in the lumen of the Golgi cisternae. These enzymes have type II protein topology with an N-terminal domain (NTD) containing a cytoplasmic tail, a transmembrane domain (TMD) and a stem region that connects the TMD to the C-terminal catalytic domain oriented toward the lumen of the Golgi (Vilcaes et al., 2011) (Fig.1A). Several molecular features within the NTD promote the retention and localization of GGTs to specific sub-Golgi compartments (Chumpen Ramirez et al., 2017; Maccioni, Quiroga, & Ferrari, 2011). The NTD additionally mediates the association and organization of GGTs into distinct multienzyme complexes (Maccioni, Quiroga, & Spessott, 2011). How the relative presence of these GGTs influence the localization of their clients is not fully elucidated in the field. The highly conserved Golgi phosphoprotein 3 (GOLPH3) is considered the first Golgi resident oncogene protein. GOLPH3 is overexpressed in many human tumors, correlating with poorer patient survival (Li et al., 2012; Wang et al., 2014; Xue et al., 2014; Zeng et al., 2012). Recent findings from our laboratory show that the physical association between ST3Gal-II and β3GalT-IV GGTs (Fig.1B) through their NTDs is mediated by GOLPH3 (Ruggiero et al., 2022). While the retention of these two enzymes at the Golgi complex does not require GOLPH3, other GGTs bind GOLPH3 via their cytoplasmic tails and this interaction influences their distribution within the Golgi as well as their protein levels through regulated lysosomal degradation (Rizzo et al., 2021). The GGT ST8Sia-I forms a ternary complex at the Golgi with the upstream GGTs (grey box in Fig.1B) thus facilitating the synthesis of b-series gangliosides (yellow box in Fig.1B). Through this association, ST8Sia-I modulates the sub-Golgi localization of the other GGTs in the complex (Uliana, Crespo, et al., 2006). These results agree with the possibility that topological distribution along the Golgi complex of GGTs that form part of multienzyme complexes relies on the relative levels of the partners (McCormick et al., 2000; Seko & Yamashita, 2005; Spessott et al., 2012). In this scenario, given that GOLPH3 is a crucial partner in the formation of the GGT complexes, it is also possible that the presence of GGTs may affect GOLPH3 localization.

**Figure 1.**
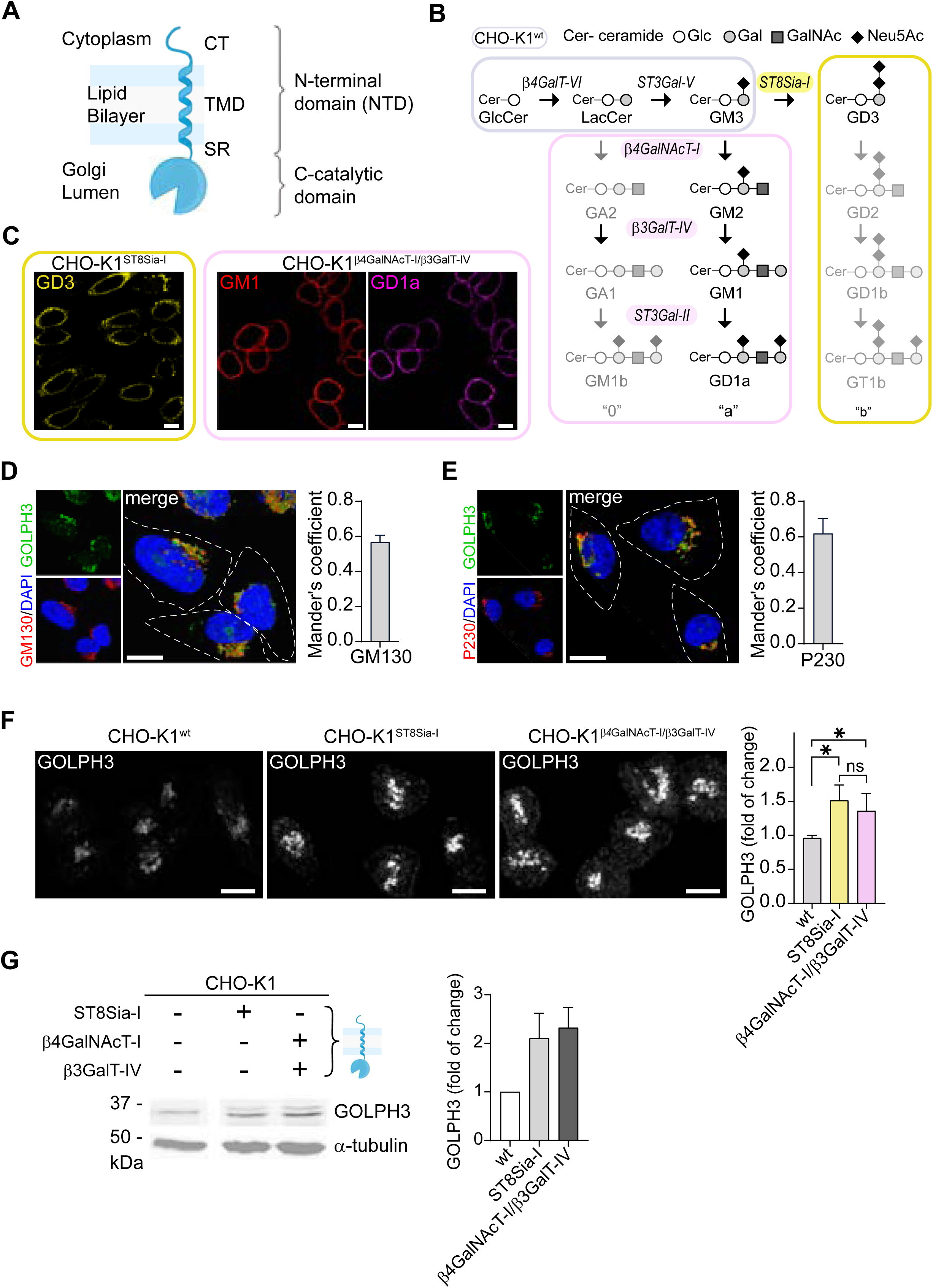
Stable expression of Ganglioside glycosyltransferases in CHO-K1 cells increase GOLPH3 expression. **(A)** Schematic representation of topology and domain organization of type II transmembrane glycosyltransferases. CT - cytoplasmic tail, TMD - Trans Membrane Domain and SR - Stem Region. **(B)** Biosynthesis pathway for 0-, a- and b-series gangliosides. Colored boxes indicate the main gangliosides expressed in CHO-K1 (grey), CHO-K1^ST8Sia-I^ (yellow) and CHO-K1^β4GalNAcT-I/β3GalT-IV^ (pink) cells. **(C)** Immunofluorescence showing GD3 (yellow) and GM1/GD1a (red/magenta) gangliosides expression in CHO-K1^ST8Sia-I^ and CHO-K1^β4GalNAcT-I/β3GalT-IV^cells respectively. **(D-E)** Representative confocal images of CHO-K1 cells stained with GOLPH3 (green), DAPI (blue) and GM130 or P230 (red; D and E respectively). Bar charts showing Manders’ coefficient (red overlapped with green). **(F-G)** Endogenous expression of GOLPH3 in CHO-K1, CHO-K1^ST8Sia-I^ and CHO-K1^β4GalNAcT-I/β3GalT-IV^ cells, analyzed by immunofluorescence **(F)** and western blot **(G)**. For immunofluorescence, statistical analysis was performed using nested one-way ANOVA with Tukey multiple comparisons. Bars represent the mean ± SD of three independent experiments. For immunoblot, statistical significance was determined using two-tailed, unpaired t test (ns: not significant). Scale bars: 10 µm.

In the present work, we demonstrate that the expression of ST8Sia-I, β3GalT-IV and ST3Gal-II enzymes increase the expression of GOLPH3 in CHO-K1 cells. Moreover, the presence of NTDs from ST8Sia-I, β3GalT-IV and ST3Gal-II GGTs, which lacks the catalytic domain, are sufficient to increase the expression of GOLPH3 in a cell type-specific manner. Moreover, our results indicate that presence of the NTD of β4GalT-VI GGT strongly inhibits the levels of GOLPH3 in cancer cell lines and consequently, reduces their capacity to undergo epithelial-mesenchymal transition. These results uncover a novel inhibitory mechanism of GOLPH3 levels with potential therapeutic applications in cancer.

## Results

### Ganglioside glycosyltransferases expression modifies GOLPH3 levels

To study the influence of glycosphingolipids metabolism on GOLPH3 expression, we took advantage of the CHO-K1 cell lines already established and well-documented in our laboratory (Fig.1B) (Rodriguez-Walker et al., 2015; Uliana, Giraudo, et al., 2006; Vilcaes et al., 2011). Wild-type CHO-K1 (CHO-K1^wt^) cell line is virtually devoid of ST8Sia-I (Daniotti et al., 2000) and β4GalNAcT-I (Rosales Fritz et al., 1997) activities, in agreement with a recent RNA-Seq data in which the ST8Sia-I gene showed almost no expression (Vishwanathan et al., 2015). Consequently, the glycosphingolipids pattern of CHO-K1^wt^ cells is dominated by glucosylceramide (GlcCer) and GM3 ganglioside (Crespo et al., 2002). ST8Sia-I glycosyltransferase is a key regulatory enzyme controlling the synthesis of b- and c-series gangliosides. The stable expression of ST8Sia-I in CHO-K1 cells (CHO-K1^ST8Sia-I^) generates primarily the ganglioside GD3 (Vilcaes et al., 2011) (Fig.1B-C). On the other hand, CHO-K1 cells stably expressing β4GalNAcT-I and β3GalT-IV glycosyltransferases (CHO-K1^β4GalNAcT-I/β3GalT-IV^) synthesize the a-series gangliosides GM1 and GD1a (Fig.1B-C). Several reports show that GOLPH3 is localized at the Golgi apparatus (Tenorio et al., 2016; Xing et al., 2016). In CHO-K1 cells, the spatial distribution of GOLPH3 is also confined mainly to the Golgi cisternae where it is alongside specific markers of cis-Golgi GM130 and trans-Golgi network P230 (Fig.1D and 1E, respectively). By immunofluorescence (IF) analysis, we found that CHO-K1^ST8Sia-I^ and CHO-K1^β4GalNAcT-I/β3GalT-IV^ cells expressed significantly higher levels of GOLPH3 compared to wild-type (WT) condition (Fig.1F). The results were also confirmed by Western Blot (WB) (Fig.1E), suggesting that the presence of these GGTs increases the expression of GOLPH3 in CHO-K1 cells.

### N-terminal domain of GGTs is sufficient to modify GOLPH3 levels in CHO-K1 cells

It has been described that the NTD of GGTs (Fig.1A) is necessary and sufficient to confer Golgi localization of GGTs (Maccioni, Quiroga, & Spessott, 2011) and is involved in covalent and non-covalent associations in some glycosyltransferase complexes (Hassinen et al., 2011). In addition, we previously demonstrated that both NTDs of β3GalT-IV and ST3Gal-II glycosyltransferases are physically associated and also that GOLPH3 interacts with both enzymes (Ruggiero et al., 2022). Thus, this prompted us to ask whether the NTDs of β3GalT-IV and ST3Gal-II enzymes influence GOLPH3 expression. To test this possibility, the NTD of ST3Gal-II fused to mCherry (ST3Gal-II^(1-51)-mCherry^) (Ruggiero et al., 2015) and the fusion protein of β3GalT-IV containing amino acids 1–52 fused to YFP (β3GalT-IV^(1-52)-YFP^) (Uliana, Crespo, et al., 2006) were expressed transiently in CHO-K1^wt^ cells. As shown in Fig.2A-B, WB analysis indicated that NTD of these enzymes were sufficient to significantly increase the levels of GOLPH3 compared to control condition. Since GGT NTDs lack the catalytic domain, our results also suggest that the increase in GOLPH3 expression is independent of gangliosides synthesis. To examine this possibility both CHO-K1^ST8Sia-I^ and CHO-K1^β4GalNAcT-I/β3GalT-IV^ cells were grown for four days in the presence of 1.2 μM P4, a GlcCer synthase inhibitor. Under these culture conditions the synthesis of glycosphingolipids including gangliosides is blocked and cell membranes are essentially devoid of them, as previously described (Vilcaes et al., 2011). As expected, P4 treatment remarkably reduced the levels of GD3 as well as GM1 and GD1a expression in CHO-K1^ST8Sia-I^ and CHO-K1^β4GalNAcT-I/β3GalT-IV^ cells, respectively (Fig.2C). However, the P4 treatment had no effect on GOLPH3 protein levels in both cell lines (Fig.2E-F). Taken together, these results support the idea that the NTD of GGTs regulate GOLPH3 expression independently of the catalytic domain or the presence of gangliosides.

**Figure 2.**
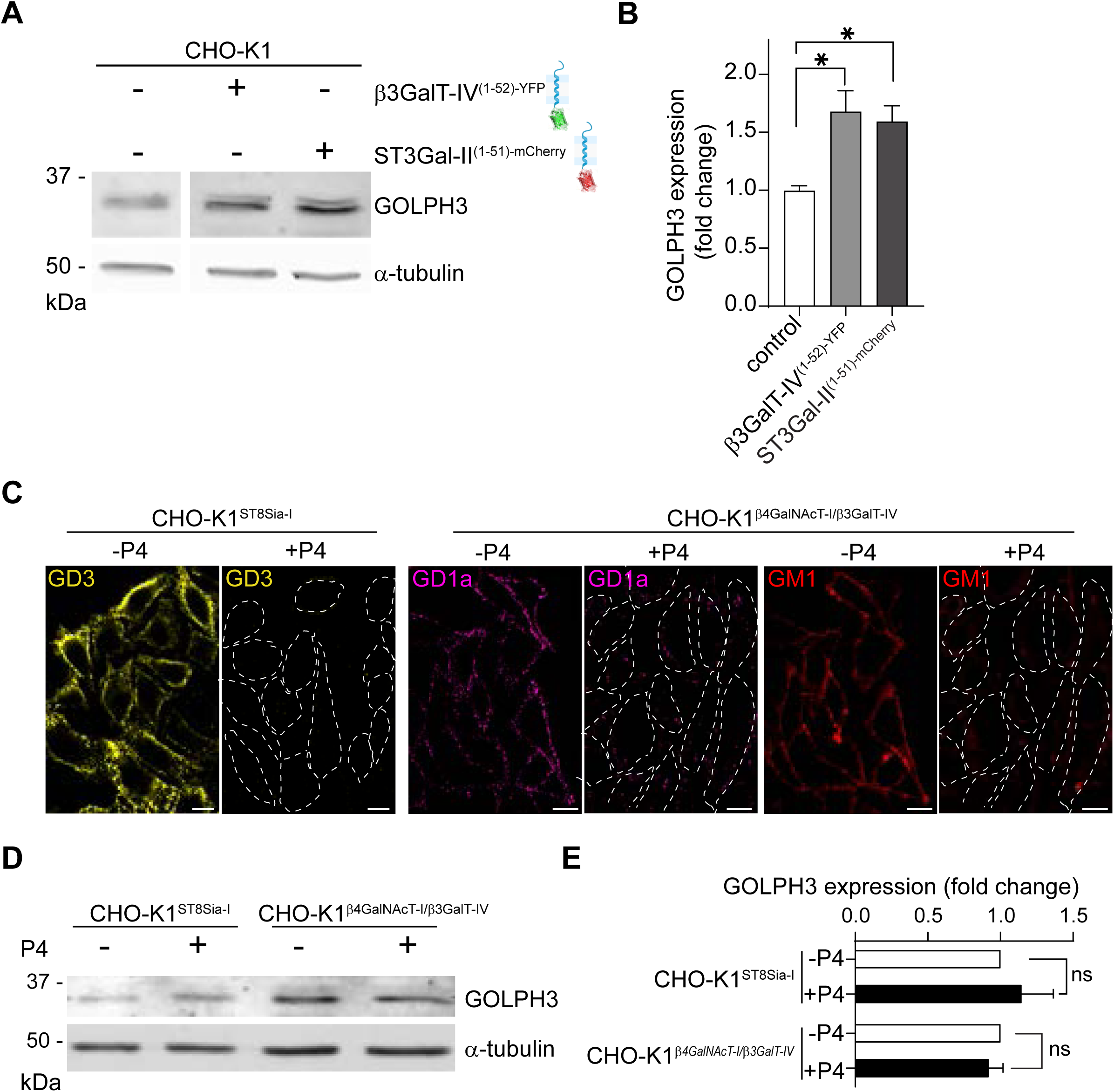
N-terminal domain of GGTs modify GOLPH3 levels independently of ganglioside synthesis in CHO-K1 cells. **(A-B)** Western Blot analysis and quantification of GOLPH3 expression in CHO-K1 cells transfected with β3Galt-IV^(1-52)-YFP^ and ST3Gal-II^(1-51)mcherry^. Data represent the mean ± SEM of three independent experiments; statistical significance was determined using one-way ANOVA. **(C)** representative images of endogenous expression of GD3 and GD1a/GM1 gangliosides in CHO-K1^ST8Sia-I^ and CHO-K1^β4GalNAcT-I/β3GalT-IV^ cells (respectively) treated with P4 (+**P4**) or vehicle (**-P4**). **(D-E)** Western Blot analysis and quantification of GOLPH3 expression from cells following the P4 treatment mentioned in C. Data represent the ± SD of three independent experiments; statistical significance was determined using two-tailed, unpaired t test (ns: not significant). Scale bars: 10 µm.

### N-terminal domain of GGTs modify GOLPH3 expression in breast cancer cells

GOLPH3 expression is elevated in various types of solid tumors and it is associated with poor overall survival in patients with breast cancer (Kuna & Field, 2019; Scott et al., 2009; Zeng et al., 2012). MDA-MB-231 and MCF7 are widely used human breast adenocarcinoma cell lines. While MDA-MB-231 are highly aggressive, invasive and have a more mesenchymal phenotype, MCF7 retain some epithelial features including hormone dependency (Theodossiou et al., 2019). Both cell lines express higher levels of GOLPH3 compared to MCF10A (Tenorio et al., 2016), a non-tumorigenic epithelial cell line widely used as *in vitro* model for studying normal breast cell function and transformation. To further explore the role of NTD domain of GGTs in GOLPH3 expression, we extended our analysis to the human breast adenocarcinomas MCF7 and MDA-MB-231 cell lines and the MCF10A human mammary epithelial cell line. To do this, ST3Gal-II^(1-51)-mCherry^, β3GalT-IV^(1-52)-YFP^ NTDs and the NTD of ST8Sia-I containing amino acids 1–57 fused to YFP (ST8Sia-I^(1-57)-YFP^) (Chumpen Ramirez et al., 2017) were expressed transiently in the three cell lines. Then, cells were fixed, and processed for localization of GOLPH3 via immunofluorescence staining. The results showed that ST8Sia-I^(1-57)-YFP^ positive cells upregulate GOLPH3 in MCF10A and MCF7 lines (Fig.3A). However, the levels of GOLPH3 in MDA-MB-231 remained unchanged in cells positive for ST8Sia-I^(1-57)-YFP^ (Fig.3A). On the other hand, the expression of β3GalT-IV^(1-52)-YFP^ (Fig.3B) or ST3Gal-II^(1-51)-mCherry^ (Fig.3C) did not modify the levels of GOLPH3 in the three cell lines tested. As mentioned before, GOLPH3 interacts with the cytoplasmic tails of several GGTs including β4GalT-V (Rizzo et al., 2021). β4GalT-V is a lactosylceramide synthase and together with β4GalT-VI are responsible for the production of lactosylceramide, a key precursor in ganglioside biosynthesis (see Fig.1B). Since β4GalT-VI forms the ternary complex β4GalT-VI/ST3Gal-V/ST8Sia-I through their N-terminal domains in CHO-K1 cells (Maccioni, Quiroga, & Spessott, 2011), it might also mimic the effect of ST8Sia-I on GOLPH3 expression. We thus examined whether the NTDs of β4GalT-V and β4GalT-VI both fused to GFP (β4GalT-V^(1-52)-GFP^ and β4GalT-VI^(1-52)-GFP^; respectively) modify GOLPH3 expression in breast cancer and mammalian epithelial cells. Unexpectedly, the expression of β4GalT-V ^(1-52)-GFP^ did not affect the expression of GOLPH3 in the three cell lines tested (Fig.4A-B). Similarly, the presence of β4GalT-VI ^(1-52)-GFP^ in MCF10A cells did not change GOLPH3 levels either. Moreover, GOLPH3 was remarkably decreased in MCF7 and MDA-MB-231 cells positive for β4GalT-VI ^(1-52)-GFP^. In summary, these findings indicate that different NTDs of GGTs can upregulate or downregulate the expression of GOLPH3 in eukaryotic cells, in a cell type-dependent manner.

**Figure 3.**
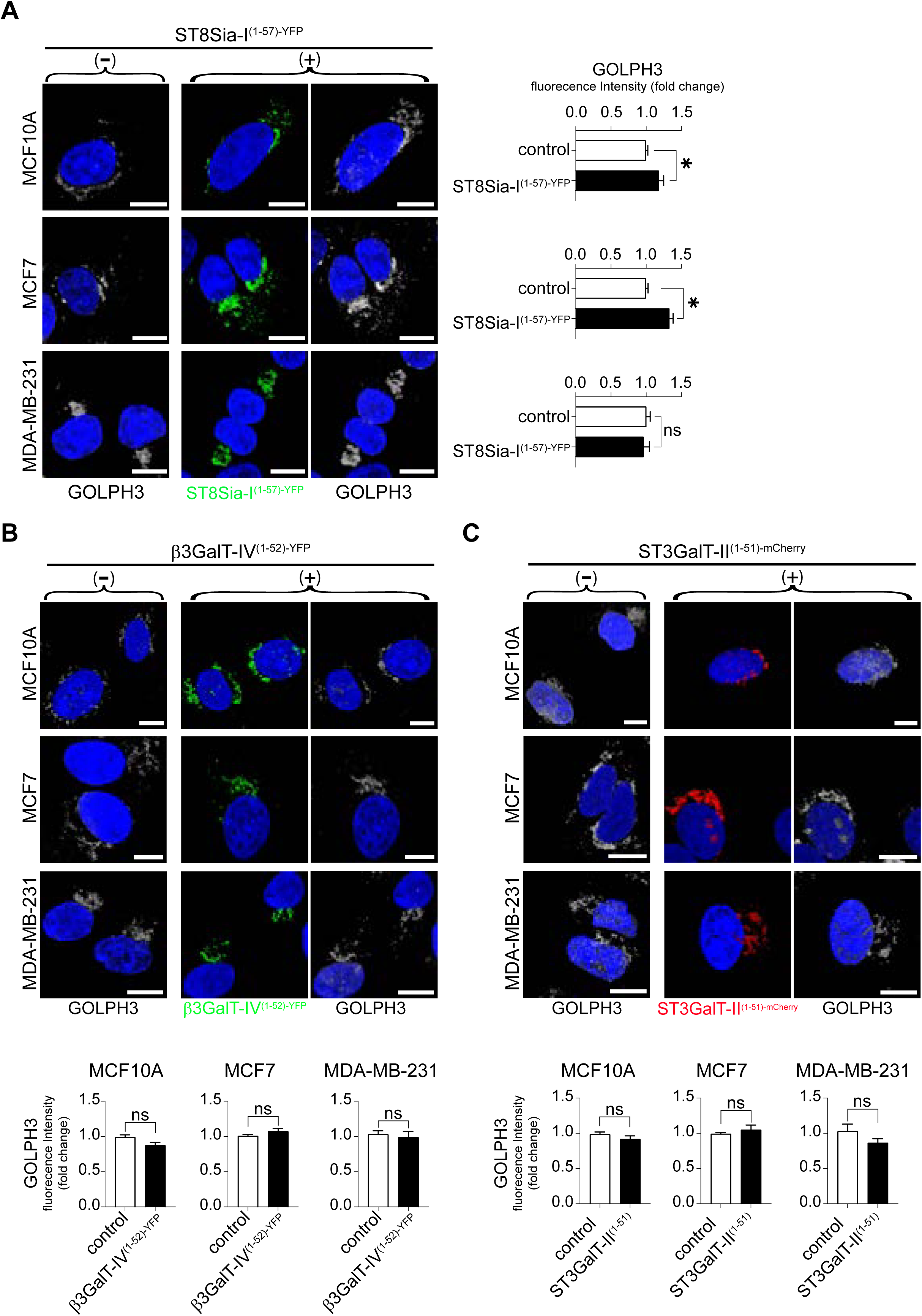
GOLPH3 expression is regulated by NTDs in a cell type-dependent manner. **(A-C)** Immunofluorescence and quantification of MCF10A, MCF7 and MDA-MB-231 cells transfected (+) with NTDs of ST8Sia-I **(A)**, β3GalT-IV **(B)**, and ST3Gal-II (C) glycosyltransferases. Cells were stained for GOLPH3 (grey) and DAPI (blue). Data represent the mean ± SD of three independent experiments, with statistical significance evaluated by nested t-test (ns: not significant). Scale bars: 10 µm.

**Figure 4.**
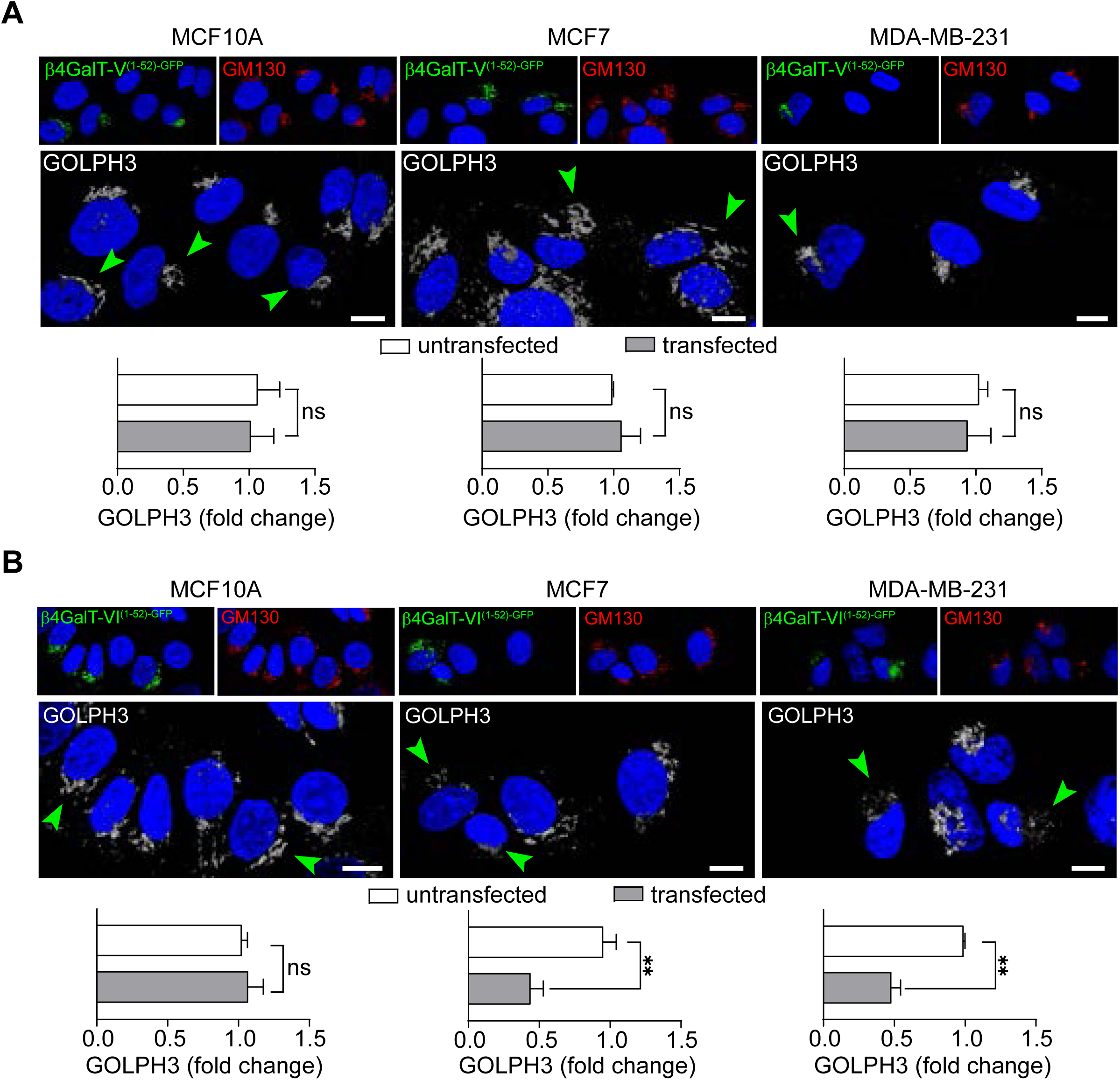
The N-terminal domain of β4GalT-V and β4GALT-VI exert opposite effects on GOLPH3 expression in breast cancer cell lines. **(A-B)** Immunofluorescence of MCF10A, MCF7 and MDA-MB-231 cells expressing the NTDs of β4GalT-V **(A)** and β4GALT-VI (B) glycosyltransferases stained for GOLPH3 (grey) and DAPI (blue). Arrowheads (green) indicate cells transfected with β4GALT-V^(1-52)-GFP^ or β4GALT-VI^(1-52)-GFP^. Comparison of GOLPH3 levels (untransfected vs transfected cells) is presented as the mean ± SD of three independent experiments, with statistical significance determined by nested t-test (ns: not significant). Scale bars: 10 µm.

### The N-terminal domain of β4GalT-VI glycosyltransferase impairs the acquisition of mesenchymal-like cancer cell features

GOLPH3 promotes the epithelial-mesenchymal transition (EMT) process, contributing to tumor growth, metastasis, and poor prognosis in several types of cancer. Also, GOLPH3 overexpression is concomitant with the upregulation of vimentin expression (Zhu et al., 2025), one of the key mesenchymal markers that is upregulated during metastasis and cancer progression (Liu et al., 2015; Usman et al., 2021). Since GOLPH3 levels are significantly downregulated in breast adenocarcinoma cells expressing the NTD of β4GalT-VI, it is then possible that β4GalT-VI may participate in the inhibition of EMT progression through the GOLPH3/Vimentin pathway. To address this, we first determined vimentin levels in cells transfected with β4GalT-VI ^(1-52)-GFP^. As shown in Fig.S1A-C, MDA-MB-231 cells expressing β4GalT-VI^(1-52)-GFP^ had decreased vimentin levels, which correlated with a diminution in GOLPH3. Since MCF7 cells do not express appreciable levels of vimentin (Liu et al., 2015), we decided to validate the effect of β4GalT-VI ^(1-52)-GFP^ in the human skin melanoma cell line SK-MEL-28, widely used for studying metastatic progression and potential therapeutic targets. Under the same experimental condition, our results confirmed a similar negative regulation on the levels of vimentin and GOLPH3 by β4GalT-VI ^(1-52)-GFP^ in this cell line (Fig.S1B-C), indicating that the regulatory effect of β4GalT-VI may extent to multiple tumor types. Collectively, these findings support the premise that the NTD of β4GalT-VI plays a role in the inhibition of EMT pathway. To evaluate this hypothesis, we induced EMT in MCF10A and MDA-MB-231 cells using TGF-β1 (Deshmukh et al., 2021) with or without the concurrent expression of β4GalT-VI ^(1-52)-GFP^ via lentiviral particles (Fig.5A). As expected, TGF-β1 treatment resulted in robust increases in vimentin and GOLPH3 levels compared to control condition in both cell lines (Fig.5B), indicating the acquisition of mesenchymal-like features. In MCF10A cells, the presence of β4GalT-VI ^(1-52)-GFP^ partially impaired the expression of vimentin and GOLPH3 in response to TGF-β1 treatment. Furthermore, β4GalT-VI ^(1-52)-GFP^ expression led to a ∼10-fold reduction in vimentin and GOLPH3 levels compared to uninfected MDA-MB-231 cells also treated with TGF-β1 (Fig.5B). Finally, we tested whether expression of β4GalT-VI NTD has preventive potential, thus rendering the cells impervious to EMT induction, by transducing MDA-MB-231 cells before TGF-β1 addition (Fig.5C). The results show that the presence of β4GalT-VI ^(1-52)-GFP^ before exposure to TGF-β1 completely abolished the increase of both GOLPH3 and vimentin (Fig.5D). Our results strongly indicate that the NTD of β4GalT-VI prevents the acquisition of EMT features, uncovering a novel potential therapeutic target for cancer prevention and treatment.

**Figure 5.**
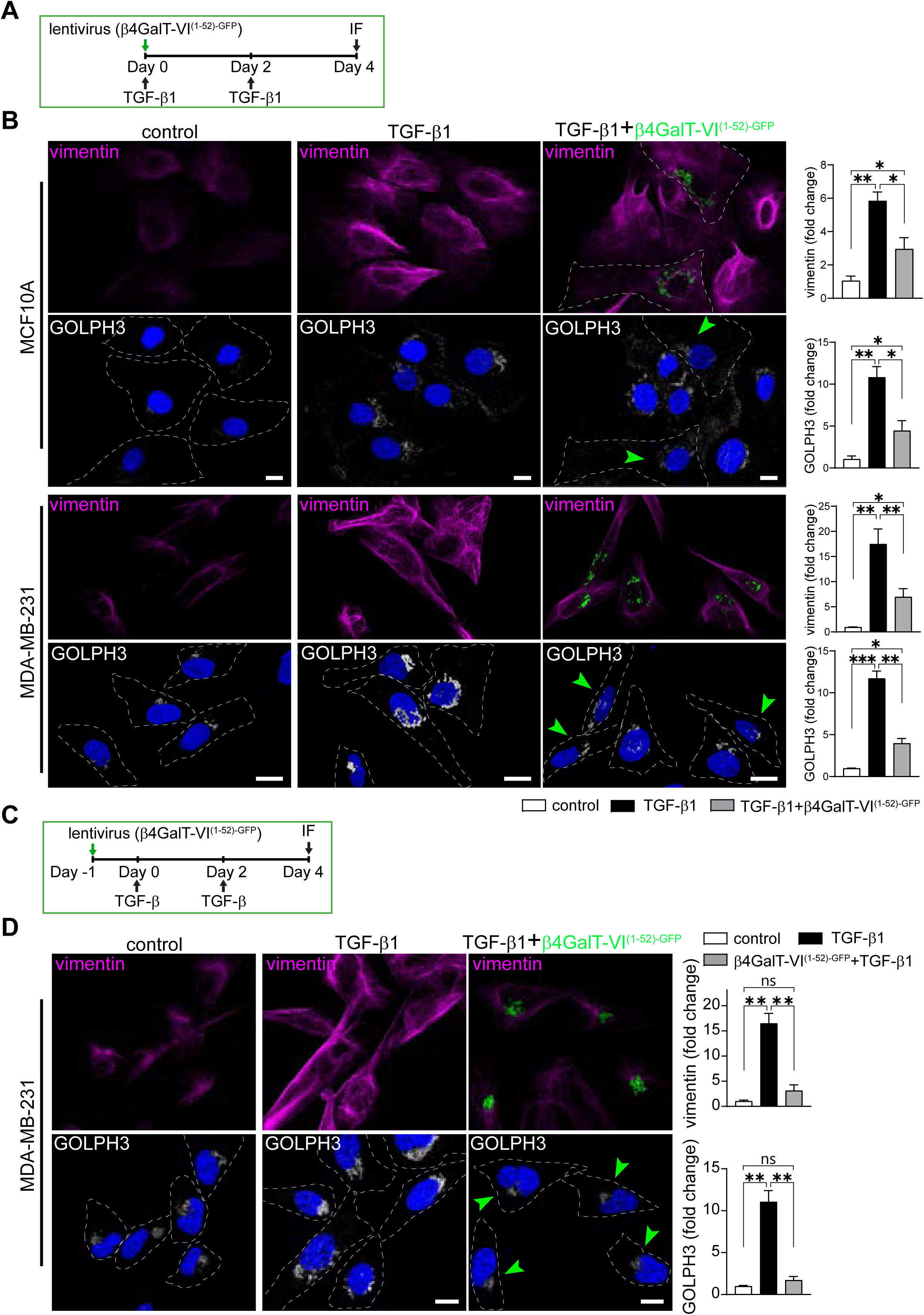
β4GalT-VI N-terminal domain prevents the acquisition of EMT features. **(A)** Schematic representation of the experimental procedure performed in B. **(B)** TGF-β1 treatment was achieved simultaneously with the lentiviral transduction of β4GALT-VI^(1-52)-GFP^. Representative confocal images of MCF10A (top) and MDA-MB-231 (bottom) cells treated with TGF-β1, TGF-β1+lentiviral particles of β4GALT-VI^(1-52)-GFP^ or without TGF-β1 treatment (control). **(C)** Schematic representation of the experimental procedure performed in D. **(D)** Lentiviral transduction of β4GALT-VI^(1-52)-GFP^ was achieved 24 h before TGF-β1 treatment. Representative confocal images of MDA-MB-231 cells treated with TGF-β1, TGF-β1+lentiviral particles of β4GALT-VI^(1-52)-GFP^ or without TGF-β1 treatment (control). Arrowheads (green) indicate cells infected with β4GALT-VI^(1-52)-GFP^. Comparison of GOLPH3 (gray) and vimentin (magenta) levels was performed using nested one-way ANOVA with Tukey multiple comparisons. Bars represent the mean ± SD of three independent experiments. Nuclei are stained with DAPI (blue). Scale bars: 10 µm.

### In silico analysis predicts different interaction surfaces between GOLPH3 and the cytoplasmic tails of β4-galactosyltransferases V and VI

Biochemical assays and bioinformatic analyses proposed that GOLPH3 interacts with the short cytoplasmic tails of numerous Golgi residents through its membrane-proximal polybasic stretches (Welch et al., 2021). Cytoplasmic tail (CT) of β4GalT-V and β4GalT-VI share a similarity of 60%. While the two arginine-rich regions in β4GalT-V CT contribute to its net positive charge, these regions are less present in β4GalT-VI CT (Fig.6A). These observations, together with the electrostatic charge distribution of GOLPH3 (face-down view from the membrane plane, Fig.6B), suggest that different regions in GOLPH3 may mediate the interaction with the CTs of β4-galactosyltransferases. We therefore used ClusPro server (https://cluspro.org) to perform computational docking of GOLPH3/β4GalT-V and GOLPH3/β4GalT-VI pairs. β4GalT-V CT was predicted to mainly interact with GOLPH3 via the electronegative region (Fig.6C), consistent with previous reports (Rizzo et al., 2021; Welch et al., 2021)). On the contrary, the CT of β4GalT-VI adopted a different orientation, binding to GOLPH3 near the phosphatidylinositol 4-phosphate (PI4P) binding pocket and the hydrophobic β-hairpin motif. In particular, β4GalT-VI R6 mediates electrostatic interactions with GOLPH3 E175 which is next to R174 residue, key in binding PI4P and thus influencing GOLPH3 localization to the Golgi (Dippold et al., 2009; Wood et al., 2009) (Fig.6C). For GOLPH3 R174, *in silico* analysis predicted hydrogen bonds and van der Waals (VW) contacts with M1, S2, V3, L4 and R5 residues from β4GalT-VI CT. A detailed recent study showed that the interactions between residues E159 and R13; D247 and R9/R12; D258/D262 and R2/R4 formed as a key binding hub of GOLPH3 and β4GalT-V complex (Theodoropoulou et al., 2025). Based on this information, we performed a ClusPro Biased Global Docking of the GOLPH3/β4GalT-V and GOLPH3/β4GalT-VI pairs, guided by the key binding hub identified above (Fig.6D). The resulting models were highly consistent with the blind docking condition. β4GalT-V CT was predicted to contact GOLPH3 via the charged surface (Fig.6D) while interaction of the CT of β4GalT-VI occurred adjacent to the PI4P binding pocket and the hydrophobic β-hairpin motif of GOLPH3. To further test the specificity of the predicted interactions between GOLPH3 and β4GalT-VI CT, we carried out a new ClusPro Global Docking but this time we performed the residue changes R2S and R4L in the CT of β4GalT-V to mimic those of β4GalT-VI CT. Under these conditions, the orientation of β4GalT-V CT^(R2S and R4L)^ became similar to that of β4GalT-VI CT (Fig.6E). Moreover, when the opposite switch was performed (S2R), β4GalT-VI CT acquired analogous orientation to β4GalT-V CT. These *in silico* results support the premise that β4GalT-VI CT^(R2S)^ interferes with GOLPH3 binding to PI4P. Previous work showed that hydrophilic residues placed at the tip of the hydrophobic β-hairpin of GOLPH3 (L195E/L196E) interfere with GOLPH3 binding to PI4P-containing liposomes and prevented its Golgi localization (Rahajeng et al., 2019). As mentioned above, docking revealed that the same hydrophobic β-hairpin (amino acids 190-201) of GOLPH3 interacts with β4GalT-VI CT primarily via hydrogen bonds and VW forces. In this model, the interactions L195 with R13; L196 with R13; F197 with R9, D198 with R9; M199 with R6 and R9; and T200 with R6 between GOLPH3 and β4GalT-VI CT, respectively, strongly suggest a reduction in the hydrophobicity of GOLPH3 β-hairpin preventing its insertion and stabilization at the Golgi membrane.

**Figure 6.**
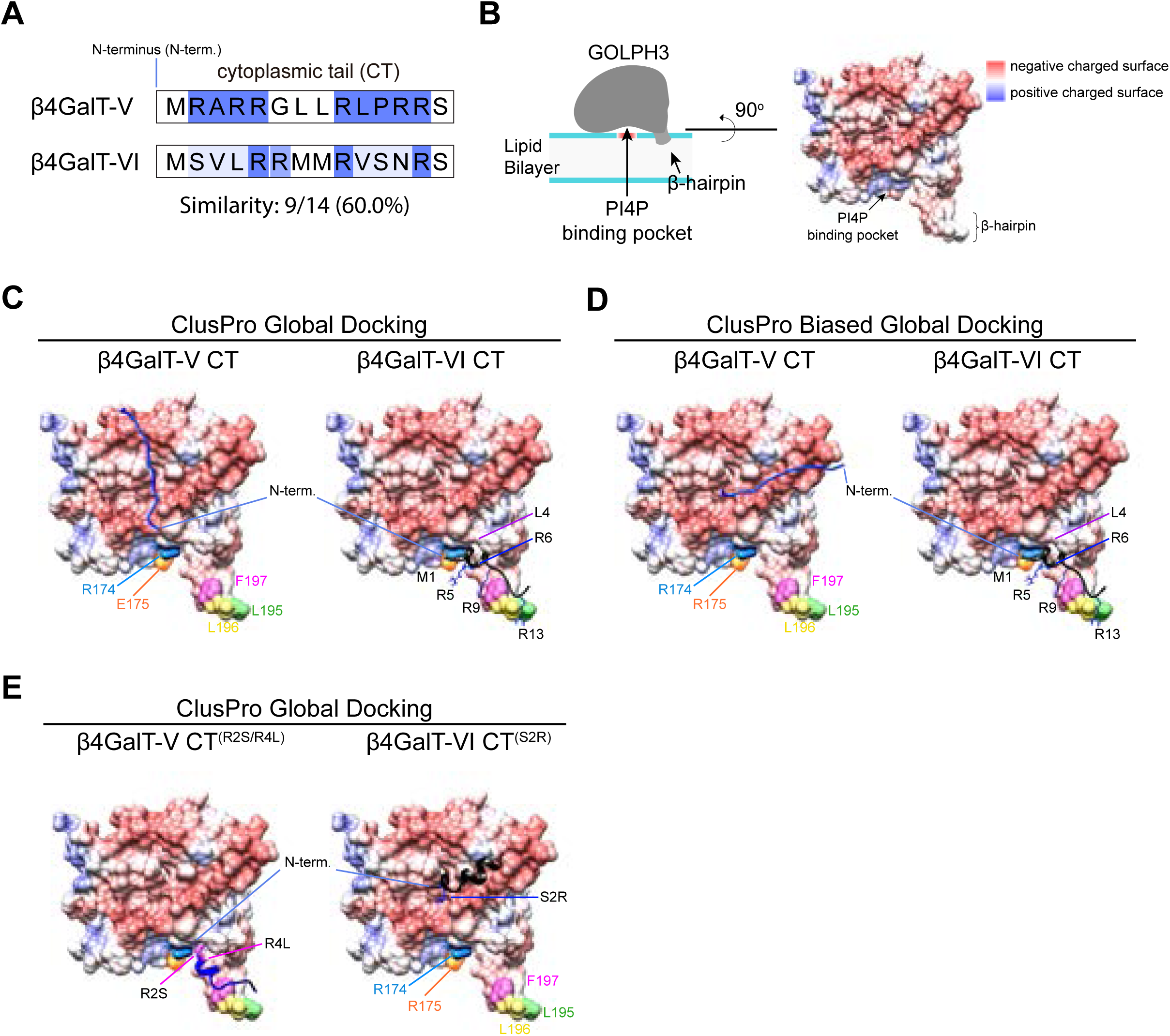
Protein–protein docking predicts differential binding of β4GalT-V and β4GalT-VI cytoplasmic tails to GOLPH3. **(A)** Homology comparison between the cytoplasmic tails of β4GalT-V and β4GalT-VI. Arginine-rich regions are shown in blue. **(B)** On the left, Scheme showing the binding of GOLPH3 to membrane (Dippold et al., 2009; Rahajeng et al., 2019; Welch et al., 2021). Arrows indicates PI4P binding pocket and hydrophobic β-hairpin motif. On the right, surface charge of GOLPH3 (generated with Chimera; red, negative; blue, positive) exposes a flat surface containing an electronegative zone (red, negative; blue, positive) (Dippold et al., 2009; Welch et al., 2021). **(C-D)** Global **(C)** and biased global **(D)** docking of β4GalT-V (blue) and β4GalT-VI (black) cytoplasmic tails with GOLPH3. Biased global docking was performed by defining D247, D258 and D262 GOLPH3 residues as attractive sites, as described previously (Theodoropoulou et al., 2025). GOLPH3 R174 and E175 residues are highlighted in cyan and orange, respectively. Side chains of putative amino acid residues in the β4GalT-V cytoplasmic tail that interact with GOLPH3 are shown. **(E)** mutants from β4GalT-V (R2S/R4L, blue) and β4GalT-VI (S2R, black) cytoplasmic tails were docked with GOLPH3 using ClusPro Global Docking. Side chains of mutated residues in β4GalT-V and β4GalT-VI cytoplasmic tails are highlighted in pink and blue, respectively.

## Discussion

The Golgi complex plays a pivotal role in the processing, sorting and transport of proteins and glycosphingolipids to different cell compartments (Iglesias-Artola et al., 2025; van Meer & Sprong, 2004). With the contributions of several laboratories, we have gained knowledge about how ganglioside glycosyltransferases (GGTs) are organized through the Golgi cisternae and how this organization is crucial for achieving the unique and precise glycosphingolipids pattern on cell surfaces. We now know that the N-terminal domain of GGTs (Fig.1A) determines their localization within sub-Golgi compartment and that the oncoprotein GOLPH3 interacts with and modulates the formation of glycosyltransferases complexes. To the extent of our knowledge, here we present a novel role of several GGTs as regulators of GOLPH3 levels in a cell-type specific manner.

CHO-K1 cells stably expressing ST8Sia-I glycosyltransferase, a key enzyme controlling b-series ganglioside biosynthesis (Fig.1B), display augmented levels of GOLPH3 respect to wild type cells which lack ST8Sia-I. Likewise, CHO-K1 cells expressing β4GalNAcT-I and β3GalT-IV glycosyltransferases, critical for the synthesis of major complex gangliosides such as GM1 and GD1a gangliosides, also show increased GOLPH3 levels. These results indicate that the presence of GGTs associated with different ganglioside synthetic pathways regulates GOLPH3 expression at the Golgi complex. Furthermore, solely the NTDs of ST3Gal-II and β3GalT-IV glycosyltransferases, which form an enzymatic complex modulated by GOLPH3, are sufficient to increase the levels of this oncoprotein. It has also been reported that GOLPH3 over-expression in HeLa cells enhances the Golgi retention and residence time of β4GalT-V, preventing its delivery to lysosomes. Collectively, our results reveal that GGTs and GOLPH3 are part of a bi-directional regulatory mechanism where they influence each other’s levels and localization.

We identified the NTD of ST8Sia-I as a likely positive regulator of GOLPH3 levels in CHO-K1 cells. Similarly, GOLPH3 levels are also increased in both human mammary MCF10A and breast carcinoma MCF7 cells expressing the NTD of ST8Sia-I. However, the presence of this NTD in the highly aggressive MDA-MB-231 cells does not change GOLPH3 expression. The transcriptional expression of the St8Sia-I gene was observed in MDA-MB-231 cells but not in MCF10A and MCF7 cells (Cazet et al., 2009). In addition, based on RNA-seq analysis from Gene Expression Omnibus (GEO) repository (Messier et al., 2016), the gene expression of ST8SIA1 in MCF10A and MCF7 cells also seems to be lower than in MDA-MB-231 cells (Fig.S2B). While GD3 ganglioside (product of St8Sia-I) is not detected in MDA-MB-231 cells (Cazet et al., 2009), the expression of GT3 ganglioside (which builds up by the action of St8Sia-I on GD3) is observed, indicating an active St8Sia-I. Under our experimental conditions, the GD3 ganglioside is also not detected in MCF10A and MCF7 cells (Fig.S2C). These data open the possibility that the expression of GOLPH3 remains unaltered in MDA-MB-231 cells after the expression of NTD from St8Sia-I because of endogenous levels of this enzyme being already plateaued leading to a ceiling effect on St8Sia-I-dependent GOLPH3 expression. We also determined that the NTDs of β3GalT-IV and ST3Gal-II do not influence the expression of GOLPH3 in MCF10A, MCF7 or MDA-MB-231 cells, contrary to the upregulation observed in CHO-K1 cells. In line with a recent study (Cavdarli et al., 2020), all three human breast cell lines have similar mRNA levels of B3GalT4 gene (GSE75168, (Messier et al., 2016)) and they also express its product, GM1 ganglioside (Fig.S2D and E). Similarly, we found that these cell lines have comparable mRNA levels for ST3Gal-II (Fig.S2F). In addition, the ST3Gal-II expression is also evidenced by the fact that MFC7 and MDA-MB-231 cells express GD1a ganglioside and SSEA4 globoside, respectively (Fig.S2G and H). These findings indicate that the bi-directional regulation between the NTD of GGTs and GOLPH3 depends on cell type, specifically on the endogenous expression pattern of different GGTs across cells, uncovering a new regulatory layer in the mechanisms that control GOLPH3 and glycosphingolipid metabolism in the Golgi complex. In line with these results, a recent study suggests that other clients of yeast GOLPH3 (Vps74) are also involved in the recruitment of Vps74 to Golgi cisternae (Lesniak et al., 2025).

The β-galactosyltransferases β4GalT-V and β4GalT-VI are responsible for the synthesis of lactosylceramide, the core structure of glycosphingolipids including gangliosides (Fig.1B). It was previously demonstrated that the NTD of β4GalT-VI physically interacts with ST3Gal-V and ST8Sia-I GGTs and that ST8Sia-I is able to promote changes in the sub-Golgi localization of β4GalT-VI. GOLPH3 also interacts with the cytoplasmic tail of several GGTs, including β4GalT-V (Rizzo et al., 2021). While β4GalT-V is expressed in various tissues, β4GalT-VI is preferentially expressed in the brain (Lo et al., 1998; Yoshihara et al., 2018). Accordingly, MCF10A, MCF7 and MDA-MB-231 cell lines have higher mRNA levels of β4GalT-V compared to the mRNA levels of β4GalT-VI (Fig.S3A). Our findings show that GOLPH3 expression remains unchanged in the different human breast cell lines positive for β4GalT-V NTD, in line with our observations showing no effects on GOLPH3 levels for NTDs of β3GalT-IV and ST3Gal-II GGTs and supporting a ceiling effect. In addition, the presence of β4GalT-VI NTD in the non-tumorigenic MCF10A epithelial cell line does not modify GOLPH3 levels as well. However, the expression of β4GalT-VI NTD in both human breast cancer cell lines, MCF7 and MDA-MB-231, promotes a significant reduction in GOLPH3 levels. Localization of GOLPH3 to Golgi membranes requires its binding to PI4P. Furthermore, the PI4P-binding capacity of GOLPH3 promotes metastasis in breast and lung cancer (Tokuda et al., 2014). To identify the molecular underpinnings of these different effects of β4GalT-V/VI CTs on GOLPH3 we performed a computational docking analysis. For β4GalT-V CT the unbiased docking predicts binding to the electronegative region of GOLPH3, agreeing with previous reports (Theodoropoulou et al., 2025; Welch et al., 2021), and validating our *in silico* approach. β4GalT-VI CT, on the other hand, was predicted to bind GOLPH3 near the PI4P-binding domain and the hydrophobic β-hairpin motif. Considering that both the PI4P binding pocket and the hydrophobic β-hairpin motif on the surface of GOLPH3 facing the membrane plane are crucial for its retention in the Golgi complex (Dippold et al., 2009; Rahajeng et al., 2019), our docking analysis thus suggests that β4GalT-VI CT may interfere with GOLPH3 ability to interact with the Golgi membrane. Further supporting these results, only exchanging a few key amino acids on the CTs of β4GalT-V and β4GalT-VI to match the other are sufficient to switch the preferred binding surface on GOLPH3.

Tumors become more invasive and malignant by undergoing the EMT process. High expression of GOLPH3 is observed in several types of cancer and is related to EMT progression and poor prognosis. Vimentin upregulation contributes to the acquisition of invasive and metastatic properties in cancer cells, in part, due to its role in regulating cell migration, cell adhesion and cytoskeletal reorganization (Ostrowska-Podhorodecka et al., 2022; Vuoriluoto et al., 2011; Wu et al., 2018). GOLPH3 stimulates EMT via the upregulation of vimentin and activation of several signaling pathways (Giansanti & Piergentili, 2022; Gong et al., 2022; Li et al., 2022; Scott et al., 2009; Sechi et al., 2020; Tan et al., 2017; Wang et al., 2020). On the contrary, the downregulation of GOLPH3 in aggressive endometrial carcinoma cells led to inhibition of vimentin expression and also inhibited the endometrial carcinoma cell invasion *in vitro* and *in vivo* by regulating EMT (Wen et al., 2019). Punicalagin, a bioactive molecule proposed as a promising chemopreventive therapy in breast cancer, also inhibits EMT by regulating GOLPH3 in breast cancer cells lines (Pan et al., 2020). Herein, we uncovered that the presence of just the NTD of β4GalT-VI promotes a reduction in vimentin and GOLPH3 levels from human melanoma and breast cancer cells. While cells concurrently treated with TGF-β1 and infected with lentiviral particles carrying the β4GalT-VI NTD partially block the upregulation of GOLPH3 and vimentin, cells infected with β4GalT-VI NTD 24 h before TGF-β1 stimulation have completely abolished increase of GOLPH3 and vimentin. Taken together, these findings strongly suggest that the CT of β4GalT-VI NTD interferes with the ability of GOLPH3 to interact with PI4P, and consequently it reduces the levels of GOLPH3 impairing its function in the acquisition of mesenchymal features.

The Golgi complex, the central trafficking hub of the cell, orchestrates the post-translational modification and sorting of proteins and lipids within the secretory pathway (Hellicar et al., 2022). Our current knowledge about the function of GGTs places them as responsible for ganglioside biosynthesis, and imbalances in glycosphingolipids metabolism cause severe metabolic disorders (Cumin et al., 2022; Daniotti et al., 2013; Morrison et al., 2025; Vo et al., 2025; Wennekes et al., 2009). Herein, we have determined a novel role of GGTs as regulators of the oncoprotein GOLPH3. The presence of pre-existing GGTs in the cell and the amino acid sequence information within the NTD of GGTs arise as new players in the mechanisms that control GOLPH3 levels in the cell. This study also opens new avenues to identify specific molecules that interfere and/or inhibit GOLPH3 pathways which will not only provide more specific tools to investigate glycolipid metabolism but also offer new therapeutic approaches with less off-target effects.

## Materials and Methods

### Cell Lines

CHO-K1, MDA MB 231, MCF7, MCF10A and SK-MEL-28 cells (ATCC, Manassas, VA, USA) were cultured at 37°C, in a humidified atmosphere with 5% CO2 in Dulbecco’s modified Eagle’s medium (DMEM) supplemented with 10% (v/v) fetal bovine serum and antibiotics (100 µg/mL penicillin and 100 µg/mL streptomycin). MCF10 cells were additionally supplemented with 0.5μg/mL hydrocortisone (H0888.10G; Sigma), 0.1µg/mL cholera toxin (C8052-1MG; Sigma) and 20ng/ml epidermal growth factor (13247-051; Invitrogen). CHO-K1 cells stable expressing ST8Sia-I and β4GalNAcT-I/β3GalT-IV were obtained as described in (Crespo et al., 2002).

### Electroporation, Transfection and Transduction

CHO-K1 cells were transfected by electroporation, 5 x 10^5^ cells were resuspended in Ingenio® Cuvettes (2mm gap) with 100 µL of electroporation mix (Ruggiero et al., 2022), containing 1µg of the corresponding plasmid and then pulsed in a BTX Electro Cell Manipulator 600 (voltage: 155 V, resistance: 186 W, capacitance: 950 µF). After electroporation, cells were seeded in a 35-mm-diameter dish and allowed for 24 h of protein expression. MDA MB 231, MCF7 and MCF10A cells were transfected with Fugene 6 transfection reagent and allowed for 24h of protein expression. Lentiviruses for β4GalT-VI ^(1-52)-GFP^ was produced in HEK293T cells by cotransfecting of pFUGW vectors and 3 packaging plasmids (pCMV-VSV-G, pMDLg/pRRE, pRSV-Rev) using Fugene 6 transfection reagent. Fresh, cleared supernatants containing lentiviruses were used for infection of MCF10A and MDA-MB-231 cells.

### Ganglioside synthesis inhibition

CHO-K1ST8Sia-I and CHO-K1β4GalNAcT-I/β3GalT-IV cells were treated with 2 μM dl-threo-phenyl-2-hexadecanoylamino-3-pyrrolidino-1-propanol (P4) for four days to reduce the glycolipid content, as described previously (Vilcaes et al., 2011). After treatment, cells were processed for immunofluorescence or western blotting. Inhibition of glycolipid synthesis was observed by immunodetection of GD3, GM1 or GD1a gangliosides.

### Plasmids

The plasmids coding for the N-terminal domains ST3Gal-II^(1-51)-mCherry^, β3GalT-IV^(1-52)-YFP^ and ST8Sia-I^(1-57)-YFP^, were constructed as described in ref. (Ruggiero et al., 2015), (Uliana, Crespo, et al., 2006) and (Chumpen Ramirez et al., 2017), respectively. The N-terminal domains of β3GalT-V and β3GalT-VI containing amino acids 1–52 were synthesized by Genscript. Then, these N-terminal domains were subcloned into the FUGW expression plasmid containing the GFP sequence after the C-terminus of the peptides.

### Antibodies

We used the following primary antibodies: polyclonal rabbit Anti-GOLPH3/MIDAS antibody (ab98023; ABCAM) (WB 1:2000), polyclonal rabbbit anti-GOLPH3 (A13121; Abclonal) (IF 1:1000), monoclonal mouse anti-GM130 (610823; BD Bioscience) (IF 1:500), monoclonal mouse anti-GD1a (#GD1a-1; DSHB) (IF 1:300), monoclonal mouse anti-GD3 (clone R24; gift from P.H.H. Lopez, CIQUIBIC-CONICET-UNC) (1:300), monoclonal mouse anti-Vimentin (AMF17b mAb; DSHB) (IF 1:1000), monoclonal mouse anti-α-tubulin (cat# T9026; Sigma-Aldrich) (WB 1:10000).

Secondary fluorochrome-conjugated antibodies were from Thermo Fisher: Alexa Fluor-488–conjugated donkey anti mouse IgG, Alexa Fluor-568–conjugated donkey anti rabbit IgG, Alexa Fluor-488–conjugated donkey anti rabbit IgG, Alexa Fluor-647-conjugated donkey anti mouse IgG. Secondary antibodies were used at a dilution 1:1000.

### Western Blot

Cells grown in 35-mm dishes for 24 h were harvested and lysed on ice for 20 min in 120 μL of lysis buffer (50 mM Tris-HCl, pH 7.2; 1% Triton X-100; 300 mM NaCl; 1 mM PMSF; cOmplete™ Mini EDTA-free protease inhibitor cocktail, Roche). Lysates were mixed with 4× Laemmli sample buffer (Bio-Rad, Hercules, CA, USA) supplemented with 5% 2-mercaptoethanol and heated at 95 °C for 10 min. Proteins were separated on 12% SDS–polyacrylamide gels under reducing conditions and transferred to nitrocellulose membranes at 350 mA for 60 min. Membranes were blocked with 5% (w/v) non-fat dry milk in PBS for 60 min and incubated overnight at 4 °C with primary antibodies. After three washes with PBS, membranes were incubated with IRDye-conjugated secondary antibodies (goat anti-mouse IgG, 800CW; goat anti-rabbit IgG, 680CW; LI-COR) diluted 1:10,000 in PBS for 60 min at room temperature. Protein bands were visualized using an Odyssey infrared imaging system (LI-COR Biotechnology, Lincoln, NE, USA). Molecular weights were estimated using calibrated protein standards run in parallel.

### Confocal Immunofluorescence Microscopy

For cell surface ganglioside labeling, cells were grown on Lab-Tek II chambered coverglass (Thermofisher), and incubated on ice for 20 min, before adding the corresponding primary antibody and/or Alexa Fluor 555-conjugated cholera toxin subunit B (C34776; Thermofisher). After a 90 min incubation, cells were washed three times with DMEM and fixed with 1% (w/v) paraformaldehyde in phosphate-buffered saline (PBS) for 5 min. For intracellular labeling, cells were fixed with 1% (w/v) paraformaldehyde in PBS and permeabilized using Saponin 0,1% in PBS. After blocking with 2% bovine serum albumin (BSA), primary antibodies were diluted in 2% BSA and incubated overnight at 4°C. Secondary antibodies were incubated 1 h at room temperature.

Confocal images were acquired using an Olympus Fluoview FV-1200 or a Zeiss LSM 980 Confocal Microscope (Carl Zeiss). Confocal z-stacks were collected with an oil immersion objective (63X; NA1.4) and 0.25 μm slices. Full z-stacks of GOLPH3, GM130, P230 and vimentin signal were taken in different regions of each coverslip/sample. Fiji 3D ImageJ Suite plugin was used for 3D segmentation and quantification of GOLPH3 levels (whole cell fluorescence intensity). For vimentin signal, whole cell fluorescence intensity quantification was performed using Z-projection tool (SUM slices) on the entire stack after background subtraction with Rolling ball algorithm. Manders’ correlation coefficient was measured using the Just Another Colocalization Plugin in ImageJ software.

### qPCR (Quantitative Real-time PCR)

SYBR Green-based qPCR was performed as we previously described (Vilcaes et al., 2020). Primers specific for human ST3Gal-II were designed and purchased from Invitrogen (Carlsbad, CA, USA). Reactions were carried out in a Rotor-Gene Q thermocycler (Qiagen, Hilden, Germany). Each 15 μl reaction contained 1 μl of cDNA template, 0.8 μM of each primer, and 7.5 μl of Real Mix (Biodynamics, Buenos Aires, Argentina). Cycling conditions were as follows: initial polymerase activation at 95 °C for 30 s, followed by 40 cycles of denaturation at 95 °C for 30 s, annealing at 60 °C for 30 s, and extension at 72 °C for 30 s. Standard curves were generated in duplicate using 1:5 serial dilutions of cDNA from MCF10A, MCF7 and MDA MB 231 cells. All samples were analyzed in triplicate. Melt curve analysis was used to confirm product specificity, and both standard curve linearity and amplification efficiency were optimized. Data were processed using Rotor-Gene Q software (Qiagen, Hilden, Germany), and relative expression levels were normalized to the geometric mean of three reference genes: PUM1, GAPDH and 18S rRNA.

### Protein-Protein docking

The crystal structure of human GOLPH3 (Wood et al., 2009) was used for docking analyses. The structure was preprocessed in UCSF Chimera v1.19 by removing water molecules and sulfate ions. The cytoplasmic tails of β4GalT-V and β4GalT-VI were obtained from the AlphaFold Protein Structure Database (https://alphafold.ebi.ac.uk, O43286 and AF-Q9UBX8-F1 respectively). Predicted full-length structures were processed in Chimera to retain only the cytosolic regions (residues 1–14). **In silico**–mutated tails were generated using the AlphaFold Server (https://alphafoldserver.com/) based on the corresponding modified amino acid sequences (CT-β4GalT-V R2S/R4L: MSALRGLLRLPRRS and CT-β4GalT-VI S2R: MRVLRRMMRVSNRS). Protein–protein docking between GOLPH3 and each cytosolic tail was performed using ClusPro (https://cluspro.org/; (Jones et al., 2022)). For biased (or guided) docking runs, residues 247, 258, and 262 of GOLPH3 were defined as attracting. The “Electrostatic-favored” models were selected for further analysis. Docking complexes were analyzed in Chimera using the Find Clashes/Contacts tool to identify residue–residue interactions. Contact lists were further processed using a custom Python script (compatible with Chimera’s internal Python 2.7 environment) to classify the interactions into hydrogen bonds, electrostatic interactions, and van der Waals contacts. Hydrogen bonds were defined by a donor–acceptor distance ≤ 3.5 Å and D–H···A angle ≥ 120°. Interaction classification was based on distance, atomic overlap, and residue charge properties.

### Bioinformatics analysis

To explore the transcriptional expression of ganglioside glycosyltransferases in MCF10A, MCF7 and MDA-MB-231 cells, we incorporated a dataset from the Gene Expression Omnibus (GEO, https://www.ncbi.nlm.nih.gov/geo/): GSE75168; (Messier et al., 2016). Dataset GSE75168 (RNA-Seq of cell lines MCF10A, MCF7 and MDA-MB-231) was used for external validation.

### Statistical Analysis

Data was plotted and analyzed using Prism software (GraphPad). For statistical analysis, data of individual cells for each independent culture was compiled into sub-columns for each experimental group and then analyzed using Nested ANOVA or Nested t test. For all experiments, when ANOVA test revealed significant effects, suitable post-tests were applied to perform multiple comparisons using the wt or control groups as reference. For simplicity the figures show asterisks for significance (* p<0.05, ** p<0.01, *** p<0.001, **** p<0.0001).

## Data availability

The data supporting the findings of this study are included in the paper and its supplemental information and will be available from the corresponding authors (Natali L. Chanaday; and A. Alejandro Vilcaes) upon request.

## Acknowledgements

We would like to thank Eve Gautreaux for her valuable feedback on the manuscript. The authors acknowledge the technical and imaging assistance of Dra. Cecilia Sampedro, Dr. Carlos Mas, Dr. Pilar Crespo and Dr. Gonzalo Quassollo from the Centro de Micro y Nanoscopía de Córdoba – CEMINCO – CONICET – Universidad Nacional de Córdoba, Córdoba, Argentina. http://ceminco.conicet.unc.edu.ar. This work was supported in part by grants from Secretaría de Ciencia y Tecnología (SECyT), Universidad Nacional de Córdoba (UNC), the University of Pennsylvania and the Margaret Q. Landenberger Research Foundation Award to N.L.C. We thank the CDB Microscopy Core (RRID SCR_022373) of the University of Pennsylvania for the use of their instruments.

## Author contributions

N. Martínez-Koteski, S. Rasino and F.M. Ruggiero: conceptualization, data curation, formal analysis, investigation, methodology, validation, visualization, and writing—original draft, review, and editing. P.H.H. Lopez and G.D. Fidelio: conceptualization, resources, and writing—review and editing. A.A. Vilcaes and N.L. Chanaday: conceptualization, data curation, formal analysis, investigation, funding acquisition, methodology, project administration, resources, supervision, visualization, and writing—original draft, review, and editing.

## Disclosures

The authors declare no competing interests.

## Supplemental data

**Figure S1.**
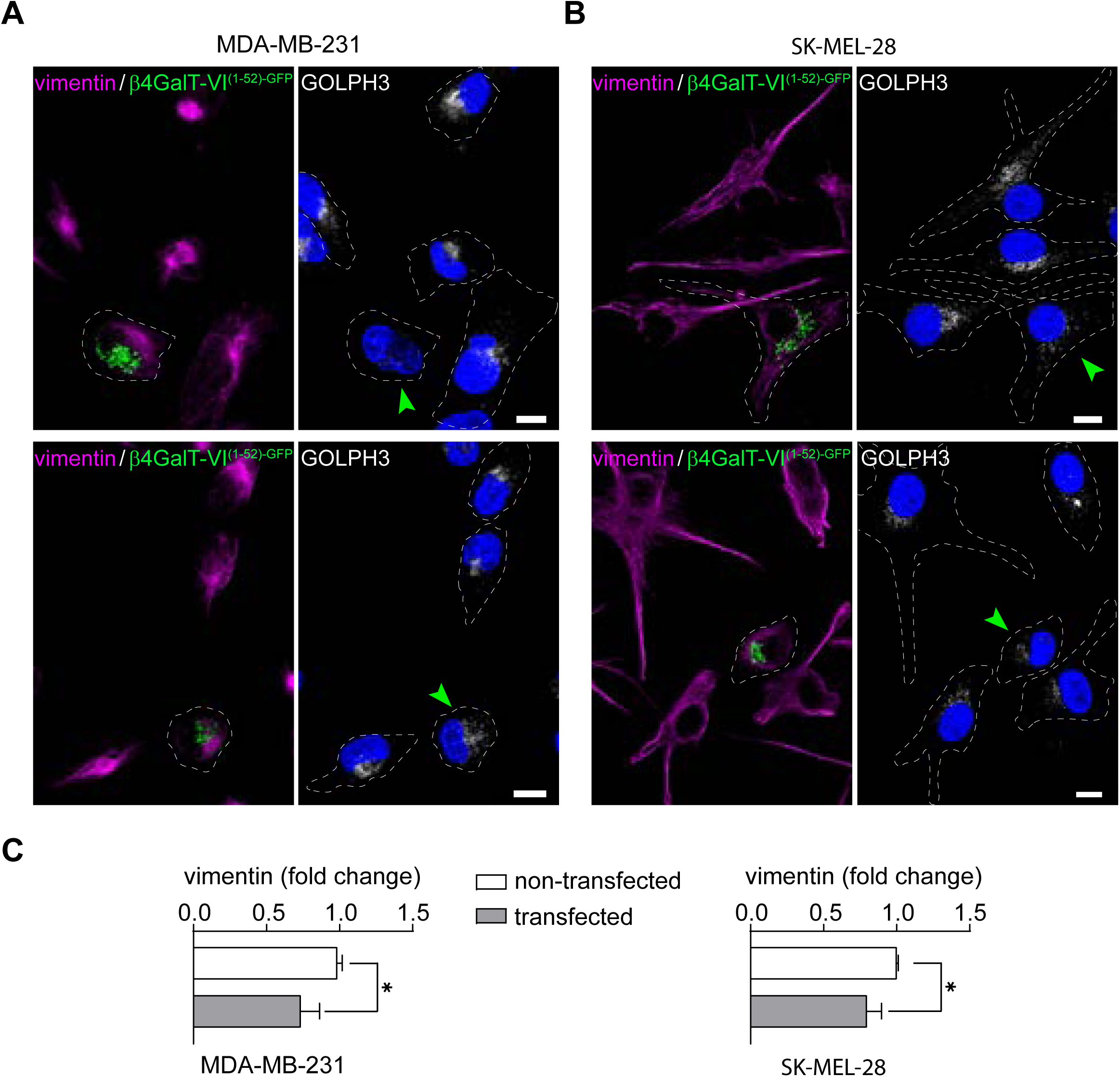
GOLPH3 and vimentin levels in MDA-MB-231 and SK-MEL-28 cells expressing β4GALT-VI*^(1-52)-GFP^*. **(A-B)** Immunofluorescence of MDA-MB-231 **(A)** and SK-MEL-28 **(B)** cells expressing β4GALT-VI^(1-52)-GFP^ stained for vimentin (magenta), GOLPH3 (grey) and DAPI (blue). Arrowheads (green) indicate cells transfected with β4GALT-VI^(1-52)-GFP^. **(C)** Comparison of GOLPH3 and vimentin levels was performed using nested t-test. Bars represent the mean ± SD of three (MDA-MB-231, bars on the left) and two (SK-MEL-28, bars on the right) independent experiments. Scale bars: 10 µm.

**Figure S2.**
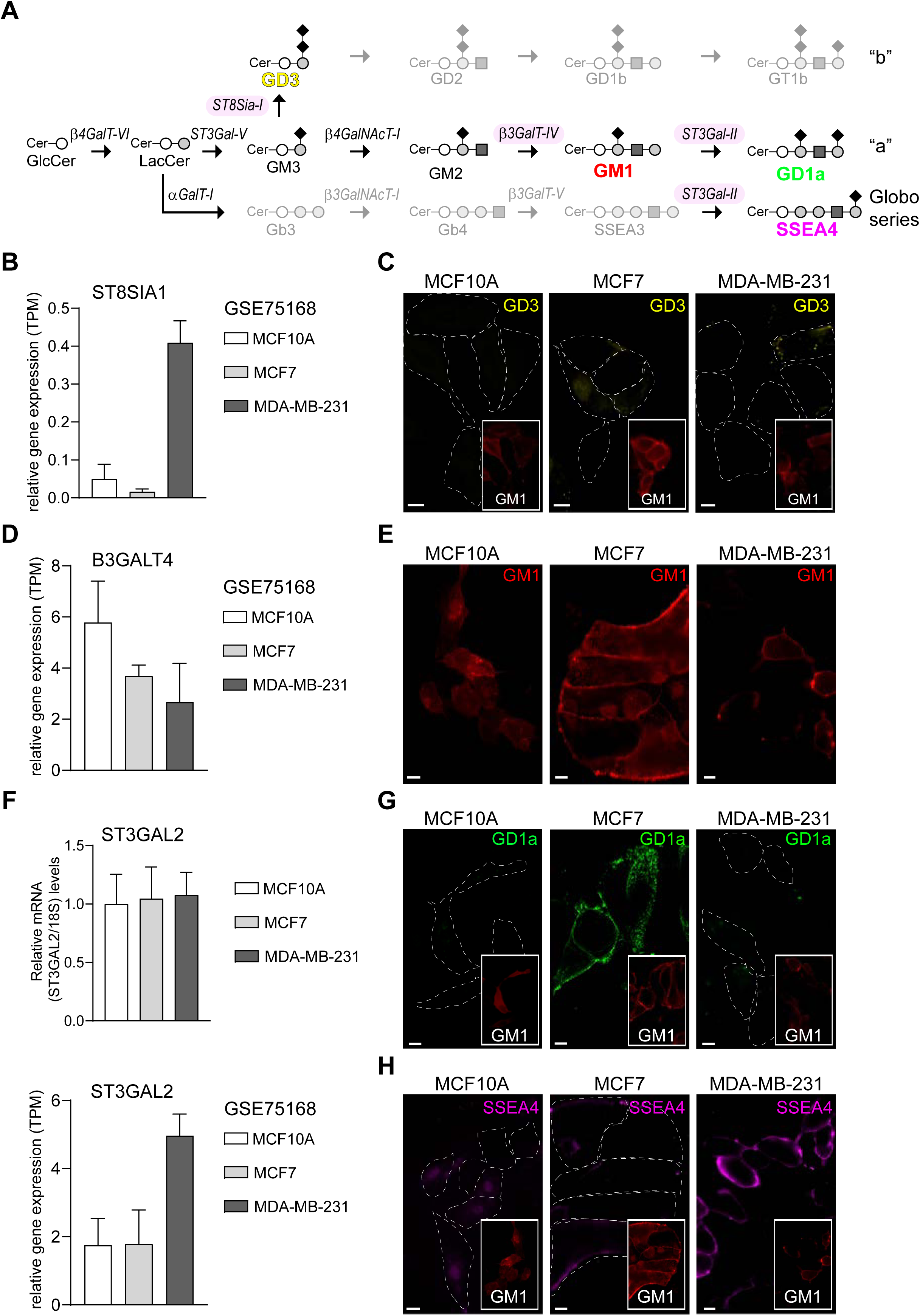
Ganglioside Glycosyltransferases gene expression profiling in human breast cell lines. **(A)** Biosynthesis pathway for Globo series and; a- and b-series gangliosides. Enzymes (highlighted in pink) that synthesize GD3 (yellow), GM1 (red), GD1a (green) and SSEA4 (magenta) are shown. **(B-D and F)** Dataset GSE75168 (Messier et al., 2016) containing RNA-Seq of MCF10A, MCF7 and MDA-MB-231 cell lines, was used for external validation. Relative gene expression of ST8SIA1 **(B)**, B3GALT4 **(D)** and ST3GAL2 (F, bottom bar charts) is shown. Each bar in the graph (averages of 3 replicates) represents the expression measurement extracted from the TPM normalized expression counts. **(C-E-G- and H)** Immunofluorescence showing GD3, GM1, GD1a and SSEA4 expression in MCF10A, MCF7 and MDA-MB-231 cells. **(F)** Top Bar charts, represent relative levels of ST3Gal-II transcripts analyzed by RT-qPCR. Total RNA was purified and reverse-transcribed from three cell lines. The expression values for RT-qPCR are given relative to the expression levels of 18S rRNA. Data represent mean ± SD from three biological replicates, each in triplicate. Scale bars: 10 µm.

**Figure S3.**
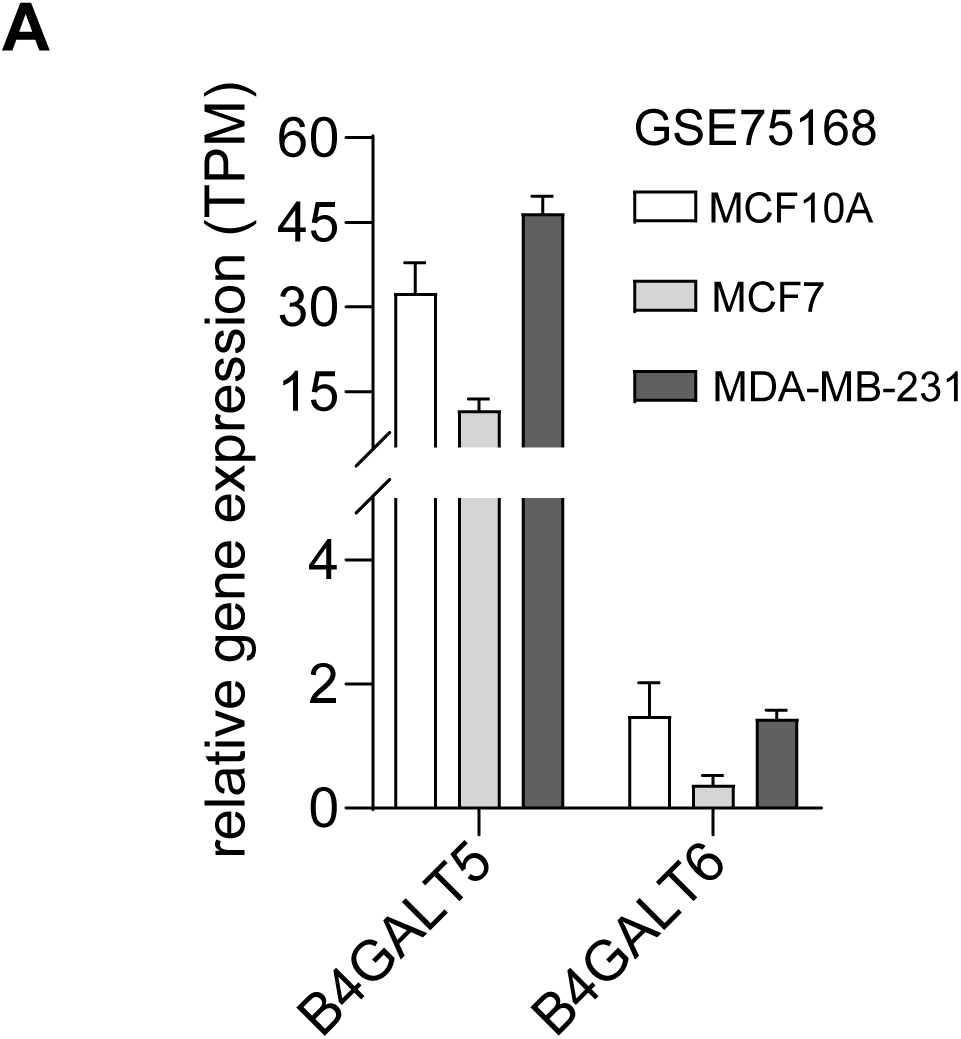
B4GALT5 and B4GALT5 gene expression profiling in human breast cell lines. **(A)** Dataset GSE75168 (Messier et al., 2016) containing RNA-Seq of MCF10A, MCF7 and MDA-MB-231 cell lines, was used for external validation. Each bar in the graph (averages of 3 replicates) represents the expression measurement extracted from the TPM normalized expression counts.

## Notes

### Competing Interest Statement

The authors have declared no competing interest.

## References

Cavdarli, S., Yamakawa, N., Clarisse, C., Aoki, K., Brysbaert, G., Le Doussal, J. M.,…Groux-Degroote, S. (2020). Profiling of. Int J Mol Sci, 21(1). 10.3390/ijms21010370

Cazet, A., Groux-Degroote, S., Teylaert, B., Kwon, K. M., Lehoux, S., Slomianny, C.,…Delannoy, P. (2009). GD3 synthase overexpression enhances proliferation and migration of MDA-MB-231 breast cancer cells. Biol Chem, 390(7), 601–609. 10.1515/BC.2009.054

Chumpen Ramirez, S., Ruggiero, F. M., Daniotti, J. L., & Valdez Taubas, J. (2017). Ganglioside glycosyltransferases are S-acylated at conserved cysteine residues involved in homodimerisation. Biochem J, 474(16), 2803–2816. 10.1042/BCJ20170124

Crespo, P. M., Zurita, A. R., & Daniotti, J. L. (2002). Effect of gangliosides on the distribution of a glycosylphosphatidylinositol-anchored protein in plasma membrane from Chinese hamster ovary-K1 cells. J Biol Chem, 277(47), 44731–44739. 10.1074/jbc.M204604200

Cumin, C., Huang, Y. L., Rossdam, C., Ruoff, F., Céspedes, S. P., Liang, C. Y.,…Jacob, F. (2022). Glycosphingolipids are mediators of cancer plasticity through independent signaling pathways. Cell Rep, 40(7), 111181. 10.1016/j.celrep.2022.111181

Daniotti, J. L., Martina, J. A., Giraudo, C. G., Zurita, A. R., & Maccioni, H. J. (2000). GM3 alpha2,8-sialyltransferase (GD3 synthase): protein characterization and sub-golgi location in CHO-K1 cells. J Neurochem, 74(4), 1711–1720.

Daniotti, J. L., Vilcaes, A. A., Torres Demichelis, V., Ruggiero, F. M., & Rodriguez-Walker, M. (2013). Glycosylation of Glycolipids in Cancer: Basis for Development of Novel Therapeutic Approaches. Front Oncol, 3, 306. 10.3389/fonc.2013.00306

Deshmukh, A. P., Vasaikar, S. V., Tomczak, K., Tripathi, S., den Hollander, P., Arslan, E.,…Mani, S. A. (2021). Identification of EMT signaling cross-talk and gene regulatory networks by single-cell RNA sequencing. Proc Natl Acad Sci U S A, 118(19). 10.1073/pnas.2102050118

Dippold, H. C., Ng, M. M., Farber-Katz, S. E., Lee, S. K., Kerr, M. L., Peterman, M. C.,…Field, S. J. (2009). GOLPH3 bridges phosphatidylinositol-4-phosphate and actomyosin to stretch and shape the Golgi to promote budding. Cell, 139(2), 337–351. 10.1016/j.cell.2009.07.052

Giansanti, M. G., & Piergentili, R. (2022). Linking GOLPH3 and Extracellular Vesicles Content-a Potential New Route in Cancer Physiopathology and a Promising Therapeutic Target is in Sight? Technol Cancer Res Treat, 21, 15330338221135724. 10.1177/15330338221135724

Gong, L. Y., Tu, T., Zhu, J., Hu, A. P., Song, J. W., Huang, J. Q.,…Chen, Y. (2022). Golgi phosphoprotein 3 induces autophagy and epithelial-mesenchymal transition to promote metastasis in colon cancer. Cell Death Discov, 8(1), 76. 10.1038/s41420-022-00864-2

Hassinen, A., Pujol, F. M., Kokkonen, N., Pieters, C., Kihlström, M., Korhonen, K., & Kellokumpu, S. (2011). Functional organization of Golgi N- and O-glycosylation pathways involves pH-dependent complex formation that is impaired in cancer cells. J Biol Chem, 286(44), 38329–38340. 10.1074/jbc.M111.277681

Hellicar, J., Stevenson, N. L., Stephens, D. J., & Lowe, M. (2022). Supply chain logistics - the role of the Golgi complex in extracellular matrix production and maintenance. J Cell Sci, 135(1). 10.1242/jcs.258879

Iglesias-Artola, J. M., Böhlig, K., Schuhmann, K., Cook, K. C., Lennartz, H. M., Schuhmacher, M.,…Nadler, A. (2025). Quantitative imaging of lipid transport in mammalian cells. Nature, 646(8084), 474–482. 10.1038/s41586-025-09432-x

Jones, G., Jindal, A., Ghani, U., Kotelnikov, S., Egbert, M., Hashemi, N.,…Kozakov, D. (2022). Elucidation of protein function using computational docking and hotspot analysis by ClusPro and FTMap. Acta Crystallogr D Struct Biol, 78(Pt 6), 690–697. 10.1107/S2059798322002741

Kuna, R. S., & Field, S. J. (2019). GOLPH3: a Golgi phosphatidylinositol(4)phosphate effector that directs vesicle trafficking and drives cancer. J Lipid Res, 60(2), 269–275. 10.1194/jlr.R088328

Lesniak, A. M., Ye, Z., & Banfield, D. K. (2025). Identification of the client-binding site on the Golgi membrane protein adaptor Vps74/yGOLPH3. iScience, 28(10), 113494. 10.1016/j.isci.2025.113494

Li, H., Guo, L., Chen, S. W., Zhao, X. H., Zhuang, S. M., Wang, L. P.,…Song, M. (2012). GOLPH3 overexpression correlates with tumor progression and poor prognosis in patients with clinically N0 oral tongue cancer. J Transl Med, 10, 168. 10.1186/1479-5876-10-168

Li, Y., Zhang, H., Long, W., Gao, M., Guo, W., & Yu, L. (2022). Inhibition of NLRP3 and Golph3 ameliorates diabetes-induced neuroinflammation. Aging (Albany NY), 14(21), 8745–8762. 10.18632/aging.204363

Liu, C. Y., Lin, H. H., Tang, M. J., & Wang, Y. K. (2015). Vimentin contributes to epithelial-mesenchymal transition cancer cell mechanics by mediating cytoskeletal organization and focal adhesion maturation. Oncotarget, 6(18), 15966–15983. 10.18632/oncotarget.3862

Lo, N. W., Shaper, J. H., Pevsner, J., & Shaper, N. L. (1998). The expanding beta 4-galactosyltransferase gene family: messages from the databanks. Glycobiology, 8(5), 517–526. 10.1093/glycob/8.5.517

Lopez, P. H., & Schnaar, R. L. (2009). Gangliosides in cell recognition and membrane protein regulation. Curr Opin Struct Biol, 19(5), 549–557. https://doi.org/S0959-440X(09)00092-X [pii] 10.1016/j.sbi.2009.06.001

Maccioni, H. J., Quiroga, R., & Ferrari, M. L. (2011). Cellular and molecular biology of glycosphingolipid glycosylation. J Neurochem, 117(4), 589–602. 10.1111/j.1471-4159.2011.07232.x

Maccioni, H. J., Quiroga, R., & Spessott, W. (2011). Organization of the synthesis of glycolipid oligosaccharides in the Golgi complex. FEBS Lett, 585(11), 1691–1698. 10.1016/j.febslet.2011.03.030

McCormick, C., Duncan, G., Goutsos, K. T., & Tufaro, F. (2000). The putative tumor suppressors EXT1 and EXT2 form a stable complex that accumulates in the Golgi apparatus and catalyzes the synthesis of heparan sulfate. Proc Natl Acad Sci U S A, 97(2), 668–673. 10.1073/pnas.97.2.668

Messier, T. L., Gordon, J. A., Boyd, J. R., Tye, C. E., Browne, G., Stein, J. L.,…Stein, G. S. (2016). Histone H3 lysine 4 acetylation and methylation dynamics define breast cancer subtypes. Oncotarget, 7(5), 5094–5109. 10.18632/oncotarget.6922

Morrison, T. A., Vigee, J., Tovar, K. A., Talley, T. A., Mujal, A. M., Kono, M.,…O’Shea, J. J. (2025). Selective requirement of glycosphingolipid synthesis for natural killer and cytotoxic T cells. Cell, 188(13), 3497–3512.e3416. 10.1016/j.cell.2025.04.007

Ostrowska-Podhorodecka, Z., Ding, I., Norouzi, M., & McCulloch, C. A. (2022). Impact of Vimentin on Regulation of Cell Signaling and Matrix Remodeling. Front Cell Dev Biol, 10, 869069. 10.3389/fcell.2022.869069

Pan, L., Duan, Y., Ma, F., & Lou, L. (2020). Punicalagin inhibits the viability, migration, invasion, and EMT by regulating GOLPH3 in breast cancer cells. J Recept Signal Transduct Res, 40(2), 173–180. 10.1080/10799893.2020.1719152

Rahajeng, J., Kuna, R. S., Makowski, S. L., Tran, T. T. T., Buschman, M. D., Li, S.,…Field, S. J. (2019). Efficient Golgi Forward Trafficking Requires GOLPH3-Driven, PI4P-Dependent Membrane Curvature. Dev Cell, 50(5), 573-585.e575. 10.1016/j.devcel.2019.05.038

Rizzo, R., Russo, D., Kurokawa, K., Sahu, P., Lombardi, B., Supino, D.,…Luini, A. (2021). Golgi maturation-dependent glycoenzyme recycling controls glycosphingolipid biosynthesis and cell growth via GOLPH3. EMBO J, 40(8), e107238. 10.15252/embj.2020107238

Rodriguez-Walker, M., Vilcaes, A. A., Garbarino-Pico, E., & Daniotti, J. L. (2015). Role of plasma-membrane-bound sialidase NEU3 in clathrin-mediated endocytosis. Biochem J, 470(1), 131–144. 10.1042/BJ20141550

Rosales Fritz, V. M., Daniotti, J. L., & Maccioni, H. J. (1997). Chinese hamster ovary cells lacking GM1 and GD1a synthesize gangliosides upon transfection with human GM2 synthase. Biochim Biophys Acta, 1354(2), 153–158. 10.1016/s0167-4781(97)00117-6

Ruggiero, F. M., Martínez-Koteski, N., Cavieres, V. A., Mardones, G. A., Fidelio, G. D., Vilcaes, A. A., & Daniotti, J. L. (2022). Golgi Phosphoprotein 3 Regulates the Physical Association of Glycolipid Glycosyltransferases. Int J Mol Sci, 23(18). 10.3390/ijms231810354

Ruggiero, F. M., Vilcaes, A. A., Iglesias-Bartolomé, R., & Daniotti, J. L. (2015). Critical role of evolutionarily conserved glycosylation at Asn211 in the intracellular trafficking and activity of sialyltransferase ST3Gal-II. Biochem J. 10.1042/BJ20150072

Scott, K. L., Kabbarah, O., Liang, M. C., Ivanova, E., Anagnostou, V., Wu, J.,…Chin, L. (2009). GOLPH3 modulates mTOR signalling and rapamycin sensitivity in cancer. Nature, 459(7250), 1085–1090. 10.1038/nature08109

Sechi, S., Frappaolo, A., Karimpour-Ghahnavieh, A., Piergentili, R., & Giansanti, M. G. (2020). Oncogenic Roles of GOLPH3 in the Physiopathology of Cancer. Int J Mol Sci, 21(3). 10.3390/ijms21030933

Seko, A., & Yamashita, K. (2005). Characterization of a novel galactose beta1,3-N-acetylglucosaminyltransferase (beta3Gn-T8): the complex formation of beta3Gn-T2 and beta3Gn-T8 enhances enzymatic activity. Glycobiology, 15(10), 943–951. 10.1093/glycob/cwi082

Spessott, W., Crespo, P. M., Daniotti, J. L., & Maccioni, H. J. (2012). Glycosyltransferase complexes improve glycolipid synthesis. FEBS Lett, 586(16), 2346–2350. 10.1016/j.febslet.2012.05.041

Tan, X., Banerjee, P., Guo, H. F., Ireland, S., Pankova, D., Ahn, Y. H.,…Kurie, J. M. (2017). Epithelial-to-mesenchymal transition drives a pro-metastatic Golgi compaction process through scaffolding protein PAQR11. J Clin Invest, 127(1), 117–131. 10.1172/JCI88736

Tenorio, M. J., Ross, B. H., Luchsinger, C., Rivera-Dictter, A., Arriagada, C., Acuña, D.,…Mardones, G. A. (2016). Distinct Biochemical Pools of Golgi Phosphoprotein 3 in the Human Breast Cancer Cell Lines MCF7 and MDA-MB-231. PLoS One, 11(4), e0154719. 10.1371/journal.pone.0154719

Theodoropoulou, A., Nasrallah, A., Abriata, L. A., Abrami, L., Talotta, F., Marcaida, M. J.,…D’Angelo, G. (2025). Molecular Regulation and Physiological Role of GOLPH3-mediated Golgi retention. bioRxiv, 2025.2006.2026.661665. 10.1101/2025.06.26.661665

Theodossiou, T. A., Ali, M., Grigalavicius, M., Grallert, B., Dillard, P., Schink, K. O.,…Berg, K. (2019). Simultaneous defeat of MCF7 and MDA-MB-231 resistances by a hypericin PDT-tamoxifen hybrid therapy. NPJ Breast Cancer, 5, 13. 10.1038/s41523-019-0108-8

Tokuda, E., Itoh, T., Hasegawa, J., Ijuin, T., Takeuchi, Y., Irino, Y.,…Takenawa, T. (2014). Phosphatidylinositol 4-phosphate in the Golgi apparatus regulates cell-cell adhesion and invasive cell migration in human breast cancer. Cancer Res, 74(11), 3054–3066. 10.1158/0008-5472.CAN-13-2441

Uliana, A. S., Crespo, P. M., Martina, J. A., Daniotti, J. L., & Maccioni, H. J. (2006). Modulation of GalT1 and SialT1 sub-Golgi localization by SialT2 expression reveals an organellar level of glycolipid synthesis control. J Biol Chem, 281(43), 32852–32860. 10.1074/jbc.M605805200

Uliana, A. S., Giraudo, C. G., & Maccioni, H. J. (2006). Cytoplasmic tails of SialT2 and GalNAcT impose their respective proximal and distal Golgi localization. Traffic, 7(5), 604–612. 10.1111/j.1600-0854.2006.00413.x

Usman, S., Waseem, N. H., Nguyen, T. K. N., Mohsin, S., Jamal, A., Teh, M. T., & Waseem, A. (2021). Vimentin Is at the Heart of Epithelial Mesenchymal Transition (EMT) Mediated Metastasis. Cancers (Basel), 13(19). 10.3390/cancers13194985

van Meer, G., & Sprong, H. (2004). Membrane lipids and vesicular traffic. Curr Opin Cell Biol, 16(4), 373–378. 10.1016/j.ceb.2004.06.004

Varki, A., Cummings, R. D., Esko, J. D., Stanley, P., Hart, G. W., Aebi, M.,…Seeberger, P. H. (2022). Essentials of Glycobiology. In. https://doi.org/NBK579905

Vilcaes, A. A., Demichelis, V. T., & Daniotti, J. L. (2011). Trans-activity of plasma membrane-associated ganglioside sialyltransferase in mammalian cells. J Biol Chem, 286(36), 31437–31446. 10.1074/jbc.M111.257196

Vilcaes, A. A., Garbarino-Pico, E., Torres Demichelis, V., & Daniotti, J. L. (2020). Ganglioside Synthesis by Plasma Membrane-Associated Sialyltransferase in Macrophages. Int J Mol Sci, 21(3). 10.3390/ijms21031063

Vishwanathan, N., Yongky, A., Johnson, K. C., Fu, H. Y., Jacob, N. M., Le, H.,…Hu, W. S. (2015). Global insights into the Chinese hamster and CHO cell transcriptomes. Biotechnol Bioeng, 112(5), 965–976. 10.1002/bit.25513

Vo, H. G., Gonzalez-Escamilla, G., Mirzac, D., Rotaru, L., Herz, D., Groppa, S., & Bindila, L. (2025). Extended coverage of human serum glycosphingolipidome by 4D-RP-LC TIMS-PASEF unravels association with Parkinson’s disease. Nat Commun, 16(1), 4567. 10.1038/s41467-025-59755-6

Vuoriluoto, K., Haugen, H., Kiviluoto, S., Mpindi, J. P., Nevo, J., Gjerdrum, C.,…Ivaska, J. (2011). Vimentin regulates EMT induction by Slug and oncogenic H-Ras and migration by governing Axl expression in breast cancer. Oncogene, 30(12), 1436–1448. 10.1038/onc.2010.509

Wang, K., Jiang, S., Huang, A., Gao, Y., Peng, B., Li, Z.,…Li, W. (2020). GOLPH3 Promotes Cancer Growth by Interacting With STIP1 and Regulating Telomerase Activity in Pancreatic Ductal Adenocarcinoma. Front Oncol, 10, 575358. 10.3389/fonc.2020.575358

Wang, Z., Jiang, B., Chen, L., Di, J., Cui, M., Liu, M.,…Su, X. (2014). GOLPH3 predicts survival of colorectal cancer patients treated with 5-fluorouracil-based adjuvant chemotherapy. J Transl Med, 12, 15. 10.1186/1479-5876-12-15

Welch, L. G., Peak-Chew, S. Y., Begum, F., Stevens, T. J., & Munro, S. (2021). GOLPH3 and GOLPH3L are broad-spectrum COPI adaptors for sorting into intra-Golgi transport vesicles. J Cell Biol, 220(10). 10.1083/jcb.202106115

Wen, Y., Tan, X., Wu, X., Wu, Q., Qin, Y., Liang, M.,…Xie, R. (2019). Golgi phosphoprotein 3 (GOLPH3) promotes endometrial carcinoma cell invasion and migration by regulating the epithelial-mesenchymal transition. Cancer Biomark, 26(1), 21–30. 10.3233/CBM-190096

Wennekes, T., van den Berg, R. J., Boot, R. G., van der Marel, G. A., Overkleeft, H. S., & Aerts, J. M. (2009). Glycosphingolipids--nature, function, and pharmacological modulation. Angew Chem Int Ed Engl, 48(47), 8848–8869. 10.1002/anie.200902620

Wood, C. S., Schmitz, K. R., Bessman, N. J., Setty, T. G., Ferguson, K. M., & Burd, C. G. (2009). PtdIns4P recognition by Vps74/GOLPH3 links PtdIns 4-kinase signaling to retrograde Golgi trafficking. J Cell Biol, 187(7), 967–975. 10.1083/jcb.200909063

Wu, S., Du, Y., Beckford, J., & Alachkar, H. (2018). Upregulation of the EMT marker vimentin is associated with poor clinical outcome in acute myeloid leukemia. J Transl Med, 16(1), 170. 10.1186/s12967-018-1539-y

Xing, M., Peterman, M. C., Davis, R. L., Oegema, K., Shiau, A. K., & Field, S. J. (2016). GOLPH3 drives cell migration by promoting Golgi reorientation and directional trafficking to the leading edge. Mol Biol Cell, 27(24), 3828–3840. 10.1091/mbc.E16-01-0005

Xue, Y., Wu, G., Liao, Y., Xiao, G., Ma, X., Zou, X.,…Huang, R. (2014). GOLPH3 is a novel marker of poor prognosis and a potential therapeutic target in human renal cell carcinoma. Br J Cancer, 110(9), 2250–2260. 10.1038/bjc.2014.124

Yoshihara, T., Satake, H., Nishie, T., Okino, N., Hatta, T., Otani, H.,…Asano, M. (2018). Lactosylceramide synthases encoded by B4galt5 and 6 genes are pivotal for neuronal generation and myelin formation in mice. PLoS Genet, 14(8), e1007545. 10.1371/journal.pgen.1007545

Zeng, Z., Lin, H., Zhao, X., Liu, G., Wang, X., Xu, R.,…Song, L. (2012). Overexpression of GOLPH3 promotes proliferation and tumorigenicity in breast cancer via suppression of the FOXO1 transcription factor. Clin Cancer Res, 18(15), 4059–4069. 10.1158/1078-0432.CCR-11-3156

Zhu, K., Fan, J., Cai, H., Zhou, C., Gong, Z., Li, Z., & Yu, J. (2025). The highly expressed. J Gastrointest Oncol, 16(2), 415–434. 10.21037/jgo-2025-193

